# Analysis of genetic requirements and nutrient availability for *Staphylococcus aureus* growth in cystic fibrosis sputum

**DOI:** 10.1101/2024.09.24.614743

**Authors:** Lauren M. Shull, Daniel J. Wolter, Dillon E. Kunkle, Katherine A. Legg, David P. Giedroc, Eric P. Skaar, Lucas R. Hoffman, Michelle L. Reniere

**Affiliations:** Department of Microbiology, University of Washington School of Medicine, Seattle, Washington, USA; Department of Pediatrics, University of Washington School of Medicine, Seattle, Washington, USA; Department of Pathology, Microbiology, and Immunology, Vanderbilt University Medical Center, Nashville, Tennessee, USA; Vanderbilt Institute for Infection, Immunology, and Inflammation, Vanderbilt University Medical Center, Nashville, Tennessee, USA; Department of Chemistry, Indiana University, Bloomington, Indiana, USA; Department of Molecular and Cellular Biochemistry, Indiana University, Bloomington, Indiana, USA; Department of Biological Sciences, Vanderbilt University, Nashville, Tennessee, USA

## Abstract

*Staphylococcus aureus* is one of the most common pathogens isolated from the lungs of people with cystic fibrosis (CF), but little is known about its ability to colonize this niche. We performed a Tn-seq screen to identify genes necessary for *S. aureus* growth in media prepared from *ex vivo* CF sputum. We identified 19 genes that were required for growth in all sputum media tested and dozens more that were required for growth in at least one sputum medium. Depleted mutants of interest included insertions in many genes important for surviving metal starvation as well as the primary regulator of cysteine metabolism *cymR*. To investigate the mechanisms by which these genes contribute to *S. aureus* growth in sputum, we quantified low-molecular-weight thiols, nutrient transition metals, and the host metal-sequestration protein calprotectin in sputum from 11 individuals with CF. In all samples, the abundance of calprotectin exceeded nutrient metal concentration, explaining the *S. aureus* requirement for metal-starvation genes. Further, all samples contain potentially toxic quantities of cysteine and sufficient glutathione to satisfy the organic sulfur requirements of *S. aureus*. Deletion of the cysteine importer genes *tcyA* and *tcyP* in the Δ*cymR* background restored growth to wild-type levels in CF sputum, suggesting that the mechanism by which *cymR* is required for growth in sputum is to prevent uncontrolled import of cysteine or cystine from this environment. Overall, this work demonstrates that calprotectin and cysteine limit *S. aureus* growth in CF sputum.

**IMPORTANCE:** *Staphylococcus aureus* is a major cause of lung infections in people with cystic fibrosis (CF). This work identifies genes required for *S. aureus* growth in this niche, which represent potential targets for anti-Staphylococcal treatments. We show that genes involved in surviving metal starvation are required for growth in CF sputum. We also found that the primary regulator of cysteine metabolism, CymR, plays a critical role in preventing cysteine intoxication during growth in CF sputum. To support these models, we analyzed sputum from 11 individuals with CF to determine concentrations of calprotectin, nutrient metals, and low-molecular-weight thiols, which have not previously been quantified together in the same samples.

## INTRODUCTION

Cystic fibrosis (CF) is a genetic disease caused by mutations in the cystic fibrosis transmembrane conductance regulator (CFTR) ion channel, which normally transports chloride ions into the extracellular space (1–3). CFTR dysfunction affects multiple organ systems, the most clinically important of which is the respiratory tract. Impaired chloride transport by airway epithelial cells leads to reduced mucus hydration, which severely impairs mucociliary clearance, a key defense against respiratory pathogens (4, 5). People with CF (pwCF) are especially vulnerable to chronic lung infection by a range of pathogens, the most common and best-studied of which are *Pseudomonas aeruginosa* and *Staphylococcus aureus* (6).

*S. aureus* is typically the earliest microbe detected in airway secretions of children with CF (7). For the past 20 years it has overtaken *P. aeruginosa* as the most prevalent bacterial species in the lungs of pwCF, with over 60% of individuals culture-positive for *S. aureus* (7). The subset of *S. aureus* infections caused by methicillin-resistant *S. aureus* (MRSA) has also been steadily increasing (7). *S. aureus* lung infections lead to heightened inflammation, frequent hospitalizations, reduced lung function which can become permanent, and lower overall survival; each of these outcomes is further worsened if the infection is caused by MRSA (8–10). These infections are typically chronic and difficult to treat with traditional antibiotics (11). Additionally, while highly-effective modulator treatments restore CFTR function, there are significant barriers to access for these treatments, including the high cost and lack of availability in many countries (12). Moreover, modulators do not eradicate bacterial infection in most pwCF (13). Despite the prevalence and importance of *S. aureus* infections in pwCF, surprisingly little is known about how *S. aureus* grows in this niche.

Modeling CF lung infection in research settings has many challenges. Animal models of CF are available in a wide range of species, including zebrafish, mice, rats, ferrets, pigs, and sheep (14). Each of these models is useful and appropriate for certain applications, but all are imperfect. For example, mice with *Cftr* mutations do not spontaneously develop lung disease, nor are they particularly susceptible to bacterial infection, making them ill-suited to model *S. aureus* lung infections (15). Only the porcine and ferret models recapitulate the lung pathology and impaired bacterial clearance observed in humans with CF, although these are both impractical for most research applications (16, 17). Airway epithelial tissue culture models have been successful in drug development applications, but are less appropriate for microbiological studies due to the absence of key aspects of infection such as mucus and immune cells (18).

An alternative to animal or tissue culture models is an *ex vivo* culture model using mucus expectorated from the lower airways, known as sputum. In the CF lung environment, *S. aureus* typically localizes within the mucus rather than on epithelial surfaces (19). This characteristic makes *S. aureus* a particularly good candidate for the *ex vivo* sputum culture model. *Ex vivo* models of infection also have the advantage of eliminating the use of animals.

With recent advancements in highly-effective CFTR modulator therapies, fewer pwCF are able to expectorate sputum. To overcome the increasing scarcity of *ex vivo* CF sputum, there has been increased interest in developing a fully synthetic *in vitro* model of growth in CF sputum. One such *in vitro* model is synthetic cystic fibrosis sputum medium (SCFM), the composition of which is based on thorough chemical analyses of CF sputum samples (20). SCFM has been thoroughly validated for modeling *P. aeruginosa* growth in CF sputum, but less data are available regarding its suitability for modeling the growth of *S. aureus* or other microbes (20–22).

In this study, we used a forward genetic screening approach to identify genes required for *S. aureus* growth in *ex vivo* CF sputum and SCFM. We also explored the heterogeneity of *S. aureus* growth in independent sputum samples and the *S. aureus* genes required for growth in each. The results of our genetic analysis were contextualized by quantifying key components that influence bacterial growth, including calprotectin, nutrient metals, and low-molecular-weight thiols, in multiple sputum samples. Overall, this work presents a comprehensive analysis of *S. aureus* growth in CF sputum and the constituents of sputum that impact this growth.

## RESULTS

### Tn-seq identifies genes required for growth in CF sputum media

To identify *S. aureus* genes required for growth in CF sputum, we performed a transposon-sequencing (Tn-seq) screen using media derived from the sputum of four pwCF. We generated a high-density transposon library containing a transposon insertion approximately every 20 base pairs in the laboratory MRSA strain JE2, derived from USA300 LAC (23, 24). Sputum media were prepared by mechanically homogenizing CF sputum at a concentration of 10% (w/v) in Staphylococcal minimal medium (SMM) with no carbon source provided (25). We confirmed the ability of each sputum medium to support the growth of *S. aureus* before proceeding with Tn-seq (data not shown). To probe both a core set of genes generally required for growth in sputum as well as heterogeneity between different samples, we inoculated the library into four individual sputum media as well as a medium composed of all four media pooled at equal volumes. As reference conditions, the library was grown in tryptic soy broth (TSB) or synthetic CF sputum medium (SCFM3), a medium developed and validated to model *Pseudomonas aeruginosa* growth in CF sputum (20, 21). The abundance of each mutant in sputum media and SCFM3 after 16 hours of growth was compared to that in TSB (Table S1). Tn-seq hits were defined as genes whose mutants were either depleted or enriched by at least two-fold in any sputum medium or in SCFM3 compared to TSB with an adjusted *p*-value of less than 0.05.

We first sought to determine the extent of overlap between the 75 to 116 hits identified in each individual sputum medium (Fig. 1A and Fig. S1A). The largest set included 39 genes that were common to all four media, suggesting that despite gross differences in appearance and other observable qualities, CF sputa from different individuals impose similar selective pressures on *S. aureus*. Of the set of 39 genes common to all four media, mutants in 19 genes were depleted, suggesting these genes are required for growth or survival in sputum (Table 1). This analysis also demonstrates that while there is a core set of genes that are important for growth in CF sputum, there is also appreciable heterogeneity among the individual sputum samples.

**Figure 1.**
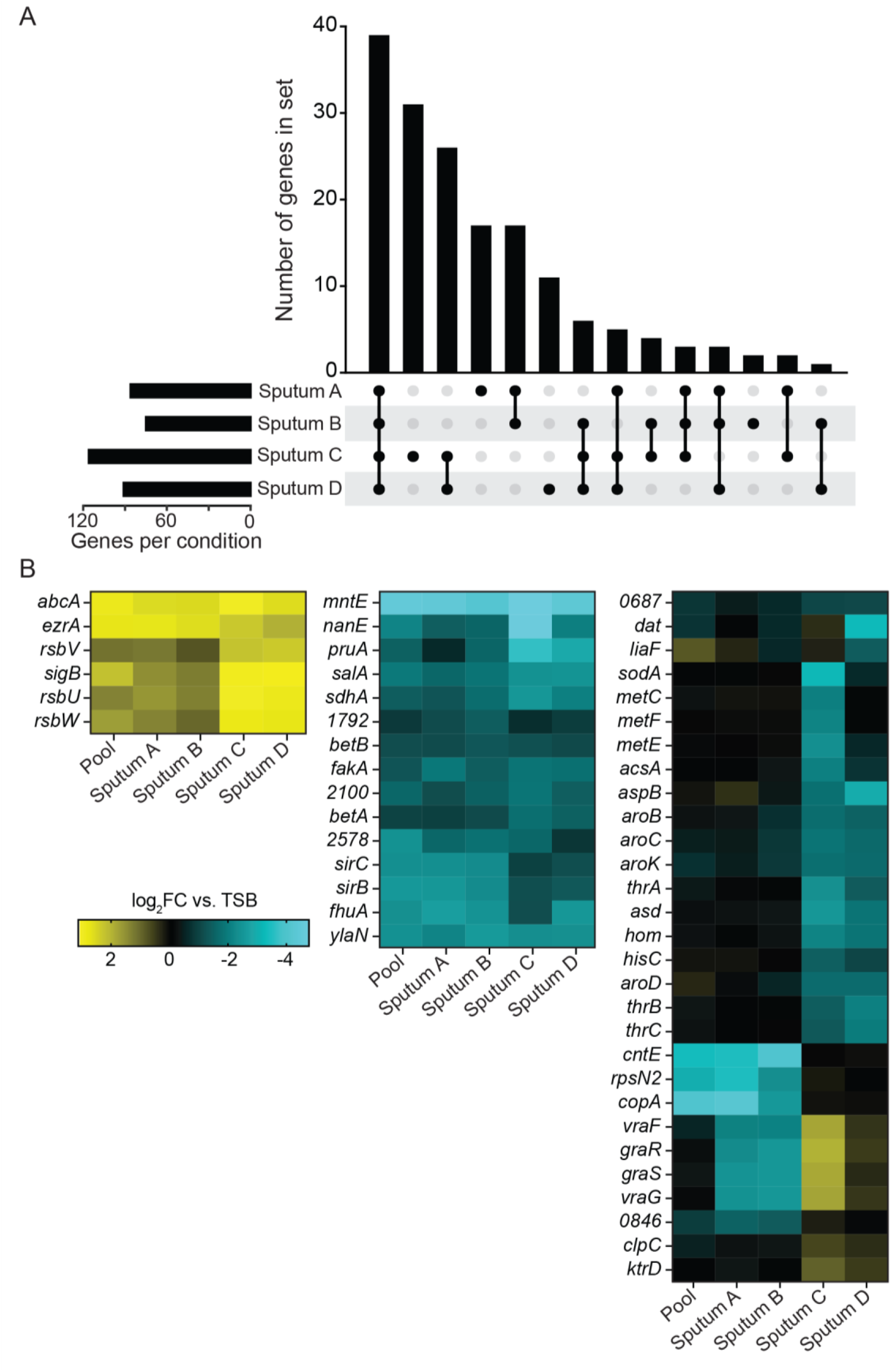
Analysis of Tn-seq results. **A)** UpSet plot showing the extent of overlap of hits between different CF sputum media. Hits were defined as genes that were significantly enriched or depleted by at least 2-fold in sputum medium compared to TSB. Significance was determined using the resampling method in TRANSIT and *p*-values were adjusted for multiple comparisons using the Benjamini-Hochberg procedure. Hits that were enriched in one or more samples but depleted in one or more other samples were not counted as overlapping. **B)** Heatmap displaying the top 50 most significant genes by one-way ANOVA with TSB as the reference condition.

**TABLE 1.** Hits significantly depleted in all four individual sputum media. Log_2_fold-change (FC) in pooled media is also indicated. Genes without annotations are identified by their SAUSA300 gene locus tags.

| Gene | Function | log<sub>2</sub>FC in sputum medium |  |  |  |  |
| --- | --- | --- | --- | --- | --- | --- |
|  |  | A | B | C | D | Pool |
| *sirB* | Iron acquisition | -2.51 | -2.26 | -1.13 | -1.32 | -2.53 |
| *nanE* | Sialic acid metabolism | -1.59 | -1.72 | -4.89 | -2.20 | -1.94 |
| *fhuA* | Iron acquisition | -2.66 | -2.44 | -1.14 | -2.47 | -2.16 |
| *ccpE* | Catabolite control protein E | -1.09 | -1.01 | -1.98 | -1.37 | -1.91 |
| *gudB* | Glutamate dehydrogenase | -1.20 | -1.29 | -1.95 | -1.45 | -1.69 |
| *ylaN* | Candidate Fur antagonist | -2.42 | -2.54 | -2.55 | -2.64 | -2.45 |
| *sdhC* | Succinate dehydrogenase complex | -1.26 | -1.49 | -2.16 | -1.77 | -1.23 |
| *sdhA* | Succinate dehydrogenase complex | -1.25 | -1.71 | -2.51 | -2.05 | -1.22 |
| *sdhB* | Succinate dehydrogenase complex | -1.19 | -1.75 | -2.20 | -1.95 | -1.61 |
| *fakA* | Fatty acid kinase | -1.89 | -1.39 | -1.80 | -1.72 | -1.23 |
| *sucB* | Succinyl-CoA synthesis | -1.39 | -1.53 | -1.92 | -1.75 | -1.39 |
| *sucA* | Succinyl-CoA synthesis | -1.14 | -1.29 | -1.21 | -1.14 | -1.26 |
| *cymR* | Cysteine metabolism regulator | -1.76 | -1.57 | -1.39 | -2.00 | -1.91 |
| *hemL* | Heme biosynthesis | -1.17 | -1.25 | -1.68 | -1.76 | -1.68 |
| *perR* | Oxidative stress response | -1.30 | -1.13 | -2.81 | -1.38 | -0.92 |
| *2100* | Putative lytic regulatory protein | -1.12 | -1.47 | -1.81 | -1.43 | -1.57 |
| *salA* | Putative chromosome partitioning protein | -1.79 | -2.01 | -2.62 | -2.60 | -2.12 |
| *mntE* | Manganese exporter | -4.42 | -4.17 | -4.77 | -4.35 | -4.36 |
| *betB* | Betaine aldehyde dehydrogenase | -1.11 | -1.28 | -1.17 | -1.06 | -1.06 |

To complement the set analysis, we next sought to cluster genes according to their importance or dispensability for growth in each sputum medium. A heatmap was generated comparing results from each individual sputum medium and pooled sputum medium to those from the TSB reference condition using an ANOVA analysis (Fig. 1B, Fig. S1B, and Table S2). This clustering strategy revealed patterns among significant genes. For example, mutants in *copA*, *rpsN2*, and *cntE* were all strongly depleted after growth in individual sputum media samples A and B as well as the pooled medium, but were neither enriched nor depleted in media from samples C and D. This suggests that a stressor present in samples A and B dominated the pooled medium even when combined with samples C and D. Mutants in the two-component system *graRS* and the primary member of its regulon, the adjacent operon *vraGF*, were all depleted in media A and B, enriched in media C and D, and neither enriched nor depleted when all four media were pooled. This pattern may indicate some opposing factors in samples A and B compared to C and D that are masked when combined. These emerging patterns highlight the power of analyzing individual sputum media, as these genes would not have been identified by only analyzing Tn-seq data from pooled sputum.

### Genes identified by Tn-seq are required for growth in CF sputum in monoculture

From the analyses described above, we selected a panel of nine genes span a variety of pathways for further study to investigate multiple mechanisms by which CF sputum imposes selective pressure on *S. aureus* (Table 2). First, genes encoding metabolic proteins were of particular interest. The transcription factor CymR is the master regulator of cysteine metabolism in *S. aureus*, suggesting a role for cysteine metabolism during growth in CF sputum (26–28).

**Table 2.** Genes selected for further study.

| Gene | Function/pathway | Condition(s) identified |
| --- | --- | --- |
| *cymR* | Cysteine metabolism regulator | Sputum A-D and pooled |
| *ylaN* | Candidate Fur antagonist | Sputum A-D and pooled |
| *nanE* | Sialic acid metabolism | Sputum A-D and pooled |
| *cntE* | Staphylopin exporter | Sputum B, D, and pooled |
| *cntA* | Staphylopin importer | None (see text) |
| *mntE* | Manganese exporter | Sputum A-D and pooled |
| *pruA* | Proline metabolism | Sputum B-D and pooled |
| *copA* | Copper efflux pump | Sputum B, D, and pooled |
| *rpsN2* | Zn<sup>2+</sup>-independent alternative S14 ribosomal protein | Sputum B, D, and pooled |

NanE and PruA belong to pathways for metabolizing sialic acid and proline, respectively (29, 30). These two nutrients are present at high concentrations in CF sputum and likely serve as nutrient sources that support bacterial growth (20, 31). MntE and CopA are manganese and copper efflux pumps, respectively, that protect cells from metal intoxication (32, 33). That these efflux pumps were required suggests that manganese and copper may be present at toxic levels in CF sputum. Next, we selected genes that suggest nutrient metals are limited in CF sputum.

Mutants lacking CntE, the exporter for the general metallophore staphylopine, are impaired for metal acquisition (34). In addition, Δ*cntE* mutants accumulate a toxic excess of staphylopine in the cytosol, which is more detrimental during zinc starvation than the lack of zinc (35). To determine whether the growth defect of the Δ*cntE* mutant was due to its zinc acquisition defect or due to staphylopine toxicity, we also evaluated a mutant deactivated for *cntA*, which was not a hit in the Tn-seq analysis. Mutants lacking *cntA* cannot import staphylopine, so these mutants exhibit the same metal acquisition defect as Δ*cntE* mutants but do not experience toxic staphylopine buildup (36). Finally, *rpsN2* encodes an alternative RpsN ribosomal protein which is zinc-independent and required during zinc starvation (37). Mutants lacking the genes-of-interest were constructed by allelic exchange or transducing a disrupting transposon from the Nebraska Transposon Mutant Library into a clean JE2 background.

To validate the Tn-seq screen, we incubated each mutant individually in TSB or a new pooled sputum medium for 12 hours and measured growth by enumerating colony forming units (CFU). To examine whether the identified genes were generalizable to different patient samples, the new pooled sputum medium combined two of the sputum samples used in the Tn-seq (Sputum C and Sputum D) with two additional samples (Sputum 7 and Sputum 8). To minimize the effects of components other than sputum on bacterial growth, these four sputum media were prepared in SCFM buffered-base rather than in SMM. All of the mutants grew similarly to the parental strain in TSB (Fig. 2A). In sputum medium, cell density of the wild-type (WT) strain increased approximately 100-fold in 12 hours, while most mutants exhibited growth defects relative to WT, three of which were statistically significant: *cymR*, *cntE*, and *rpsN2* (Fig. 2B).

**Figure 2.**
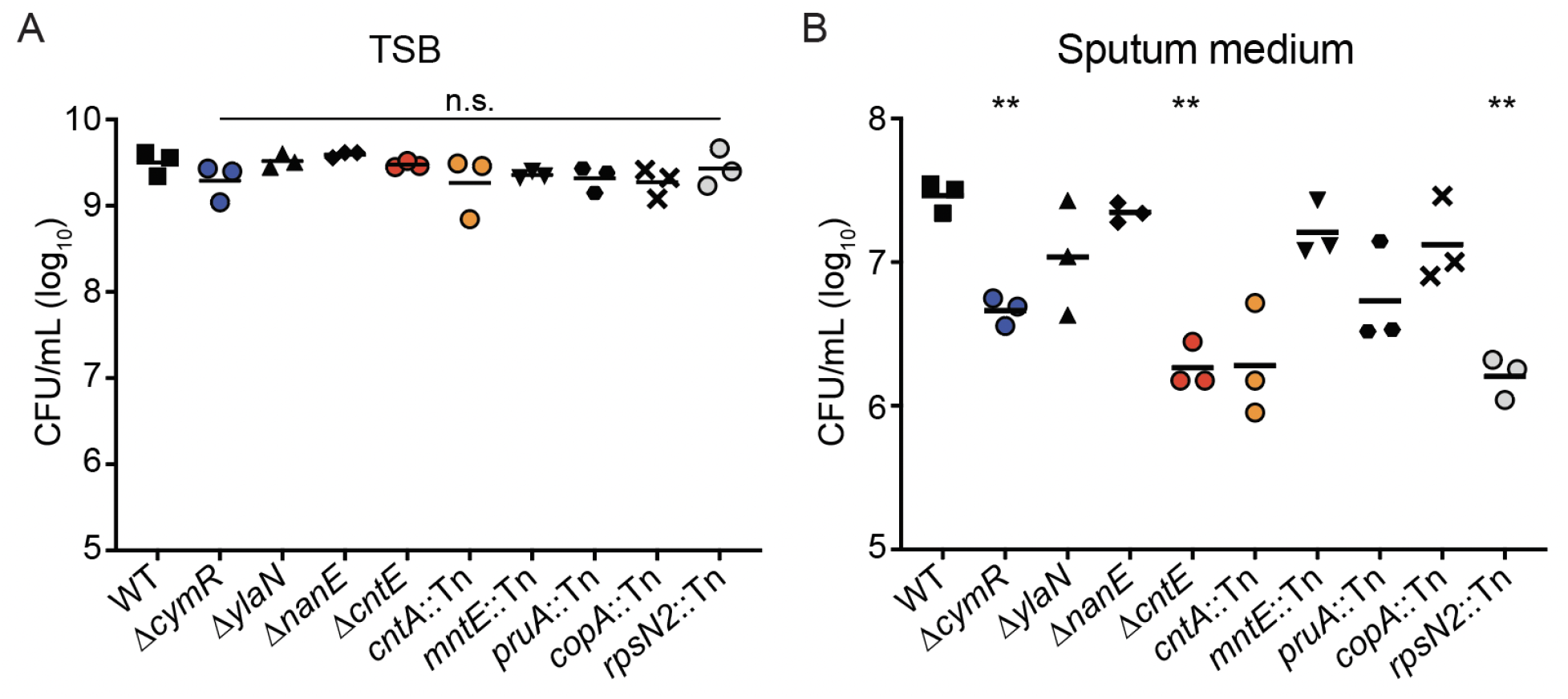
Bacterial density after 12 hours of growth. Approximately 2 x 10^5^ CFU of each strain was inoculated and incubated for 12 hours in 100 µL of **A)** TSB or **B)** pooled CF sputum media. Data represent three independent experiments combined. Two asterisks (**) indicate *p* < 0.01 as measured by Brown-Forsythe & Welch One-way ANOVA with Dunnett’s multiple comparison correction on log-transformed CFU values and n.s. indicates *p* > 0.05 compared to WT.

These results demonstrate the robustness of the Tn-seq screen, as many mutants displayed growth defects in sputum in monoculture, as well as in sputum from patients beyond those in the original screen.

### Calprotectin restricts metal availability in sputum

Host sequestration of essential nutrient transition metals has long been appreciated as a key defense against bacterial infections, in a process known as nutritional immunity (38–40). Many of the hits we identified by Tn-seq are required for surviving metal starvation, suggesting that CF sputum is also devoid of these key nutrients. Mutants that lacked *cntE* or *rpsN2*, genes known to be required to survive zinc starvation, exhibited significant growth defects in pooled CF sputum medium (41, 42). Several other genes identified by Tn-seq as required for growth in CF sputum, including *ylaN*, *fhuA*, and *sirB*, are necessary during iron limitation (43–45). In contrast, *copA*, which encodes a copper efflux pump, was important in a subset of sputum media, suggesting that copper availability may impact *S. aureus* growth more in some CF sputa than in others (33).

To investigate the importance of metal starvation during *S. aureus* growth in CF sputum, we quantified total transition metals in sputum samples from 11 pwCF using inductively coupled plasma mass spectrometry (ICP-MS). Copper levels ranged from 164 to 654 nM, with an average of 325 nM (Fig. 3). Iron was generally more abundant, with concentrations ranging from 0.57 to 3.61 µM and an average of 1.61 µM. Zinc was the most abundant transition metal, ranging from 0.61 to 6.27 µM and an average of 2.59 µM. While SCFM contains neither copper nor zinc, it does contain iron at a concentration of 3.6 µM, comparable to the concentrations we measured in sputum. Additional elements are quantified in Figure S3.

**Figure 3.**
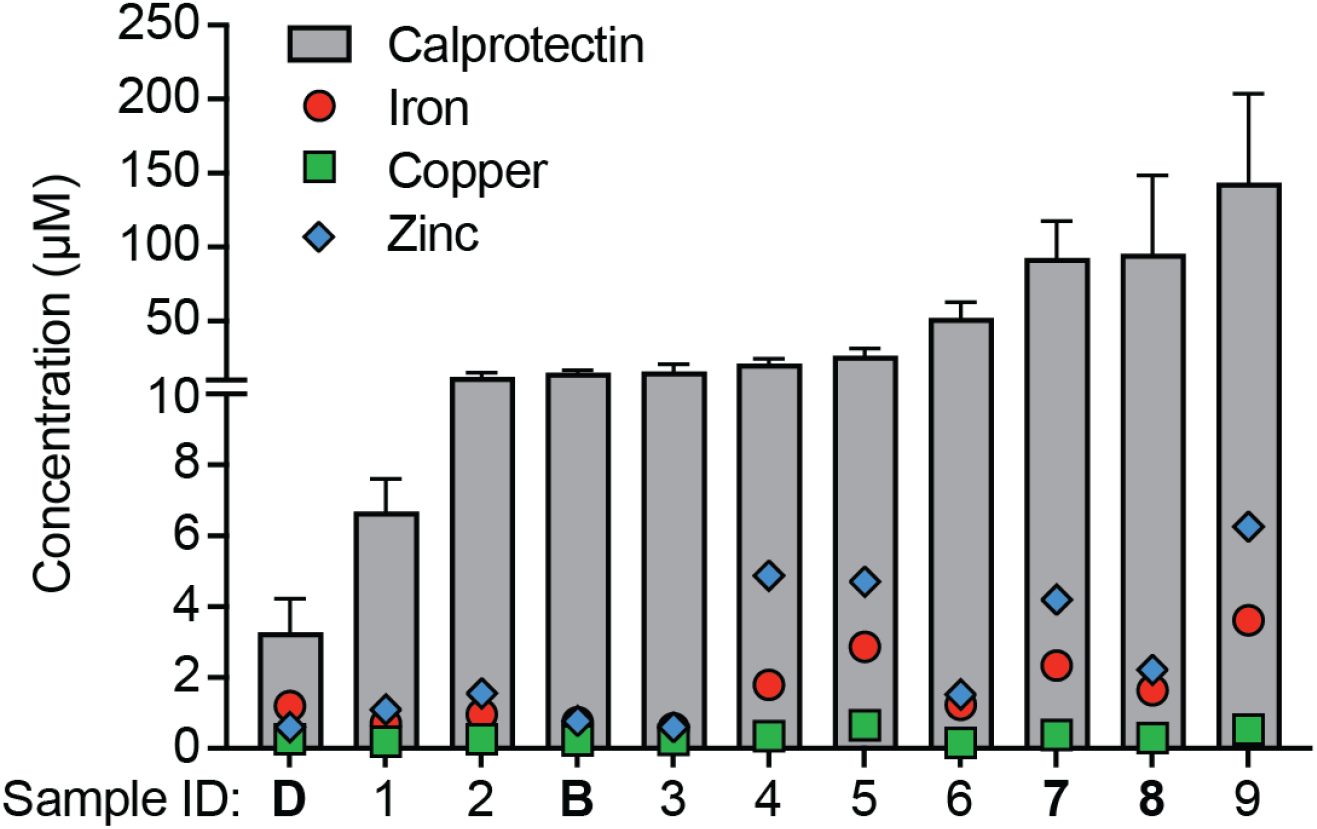
Quantification of calprotectin, iron, zinc, and copper in CF sputum. Total metals were quantified by ICP-MS while calprotectin was quantified by ELISA. Calprotectin quantifications are the mean and standard deviation (SD) of at least two independent assays, each performed in technical duplicate. The four samples used in validation medium in Fig. 2 are indicated in bold. Samples B and D were also used in Tn-seq.

A host protein that is particularly important in nutritional immunity is calprotectin, a dimer of calcium-binding proteins S100A8 and S100A9, which can bind and sequester a spectrum of transition metals with affinities in the picomolar to nanomolar range (40, 46–48). Calprotectin is primarily produced by neutrophils and thus is often abundant at sites of infection, including chronic infections, such as those occurring in the lungs of pwCF (49). As ICP-MS quantifies all metals in a sample, whether or not they are protein-bound, we contextualized metal availability and quantified calprotectin by ELISA in the same sputum samples. In all 11 cases, calprotectin was in significant excess relative to any metal we quantified (Fig. 3). These observations suggest the simple model that the mechanism by which *cntE* and *rpsN2* are required for growth in CF sputum is that mutants lacking these genes are unable to survive metal starvation imposed by calprotectin.

### Cysteine concentrations in sputum

In addition to *cntE* and *rpsN2*, *cymR* was also required for *S. aureus* growth in CF sputum, yet the importance of *cymR* in this growth niche cannot adequately be explained by calprotectin-dependent metal sequestration. CymR is a redox-sensitive transcription factor whose regulon consists of genes involved in metabolizing and importing organic sulfur sources, particularly cysteine (26–28). Deletion of *cymR* is known to sensitize *S. aureus* to excess cysteine, and thus, we hypothesized that the growth defect of the Δ*cymR* mutant in sputum may be caused by excess cysteine (26). To address this hypothesis, we quantified the low-molecular-weight (LMW) thiols cysteine and glutathione (GSH) using an isotope-dilution LC-MS workflow in the same 11 sputum samples in which we quantified metals and calprotectin. Patient samples were not stored in specific anaerobic conditions, and therefore the ratios of oxidized (cystine and glutathione disulfide, GSSG) to reduced thiols (cysteine and GSH) present in the samples at the time of expectoration could not be determined. Rather, we quantified the total amounts of each thiol after treating these samples with a reducing agent. The sputum samples contained between 11 and 296 µM GSH, higher than the published quantity of approximately 10 µM (Fig. 4) (50). This concentration of GSH alone is sufficient to fulfill the organic sulfur requirements of *S. aureus* (51). Total cysteine concentrations in CF sputum ranged from 65 to 3350 µM, and 10 of 11 samples surprisingly contained a higher concentration of cysteine than the 160 µM present in SCFM (Fig. 4) (20). It is therefore plausible that cysteine concentrations in this range could be toxic to the Δ*cymR* mutant during growth in CF sputum.

**Figure 4.**
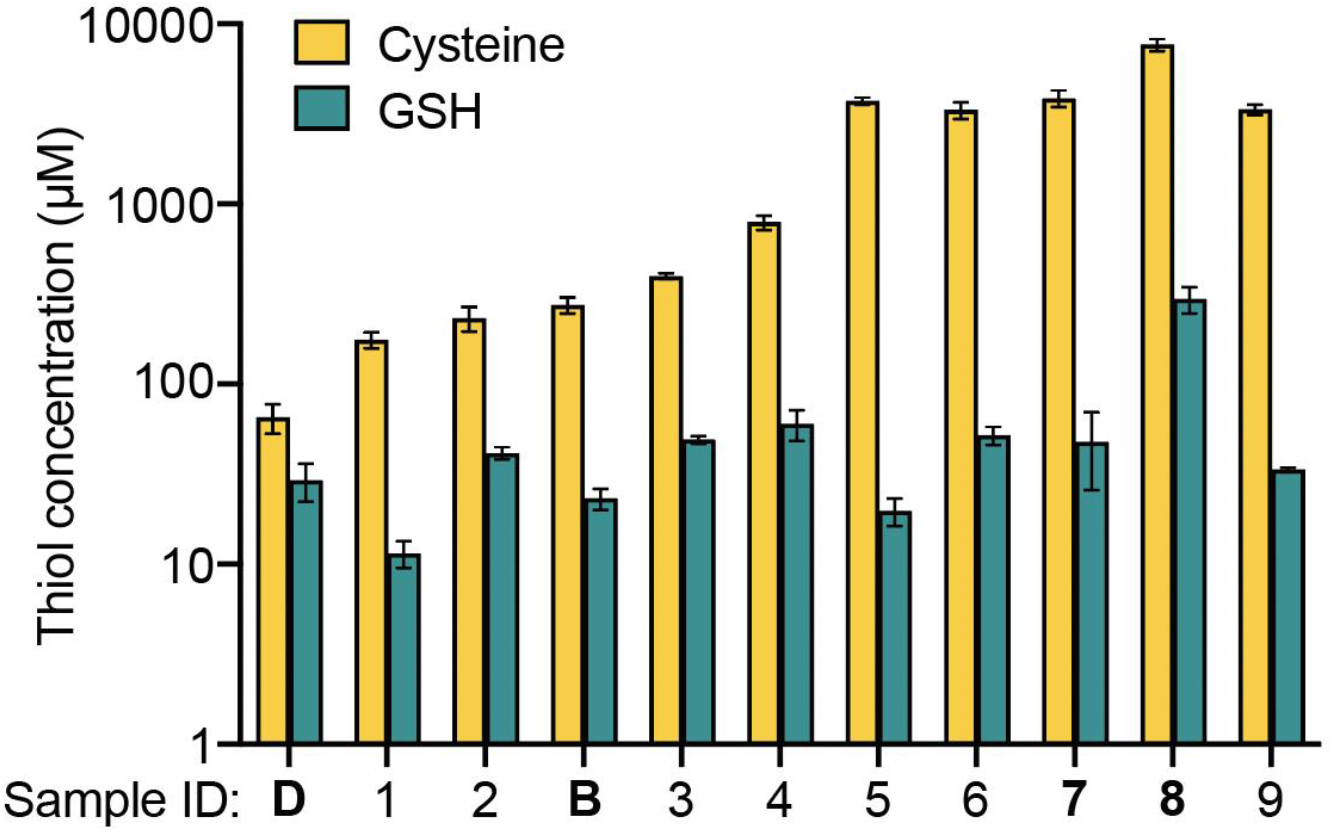
Quantification of thiols in CF sputum. The same 11 sputum samples from Figure 3 were analyzed by LC-MS for thiol abundance after treatment with a reducing agent. Data are the mean and SD of triplicate measurements. Samples used in validation medium are indicated in bold.

### Excess cysteine import is toxic during growth in CF sputum

To investigate the role of *cymR* during growth in CF sputum, we took a genetic approach and employed an *in vitro* synthetic sputum model, as manipulating the components of *ex vivo* sputum is not feasible. We used SCFM1 for these experiments, which differs from SCFM3 primarily by omitting mucin and DNA, while including all amino acids, including cysteine, at the same concentrations as SCFM3 (21). The absence of mucin renders SCFM1 optically clear, enabling bacterial growth to be measured by optical density. To test the hypothesis that the relative growth defect of the Δ*cymR* mutant in CF sputum is due to cysteine toxicity, we grew WT *S. aureus*, the Δ*cymR* mutant, and a *cymR* genetic complement in SCFM1 (20). The Δ*cymR* mutant and its complement grew similarly to WT in TSB (Fig. 5A). However, the Δ*cymR* mutant displayed reduced growth in SCFM1 compared to WT, but growth was restored when *cymR* was reintroduced on a plasmid under the control of its native promoter (Fig. 5B). Interestingly, the Δ*cymR* mutant did not exhibit a growth defect when grown in TSB supplemented with 2 mM cysteine, indicating that the observed phenotype in SCFM1 varies depending on nutritional context (Fig. 5C). The concentration of cysteine in SCFM is 160 µM, which is a very conservative estimate of cysteine availability in CF sputum when compared with our thiol profiling data (Fig. 4). It is striking that cysteine is toxic to the Δ*cymR* mutant in SCFM1 even at this low concentration, but not at the high concentration of 2 mM in TSB.

**Figure 5.**
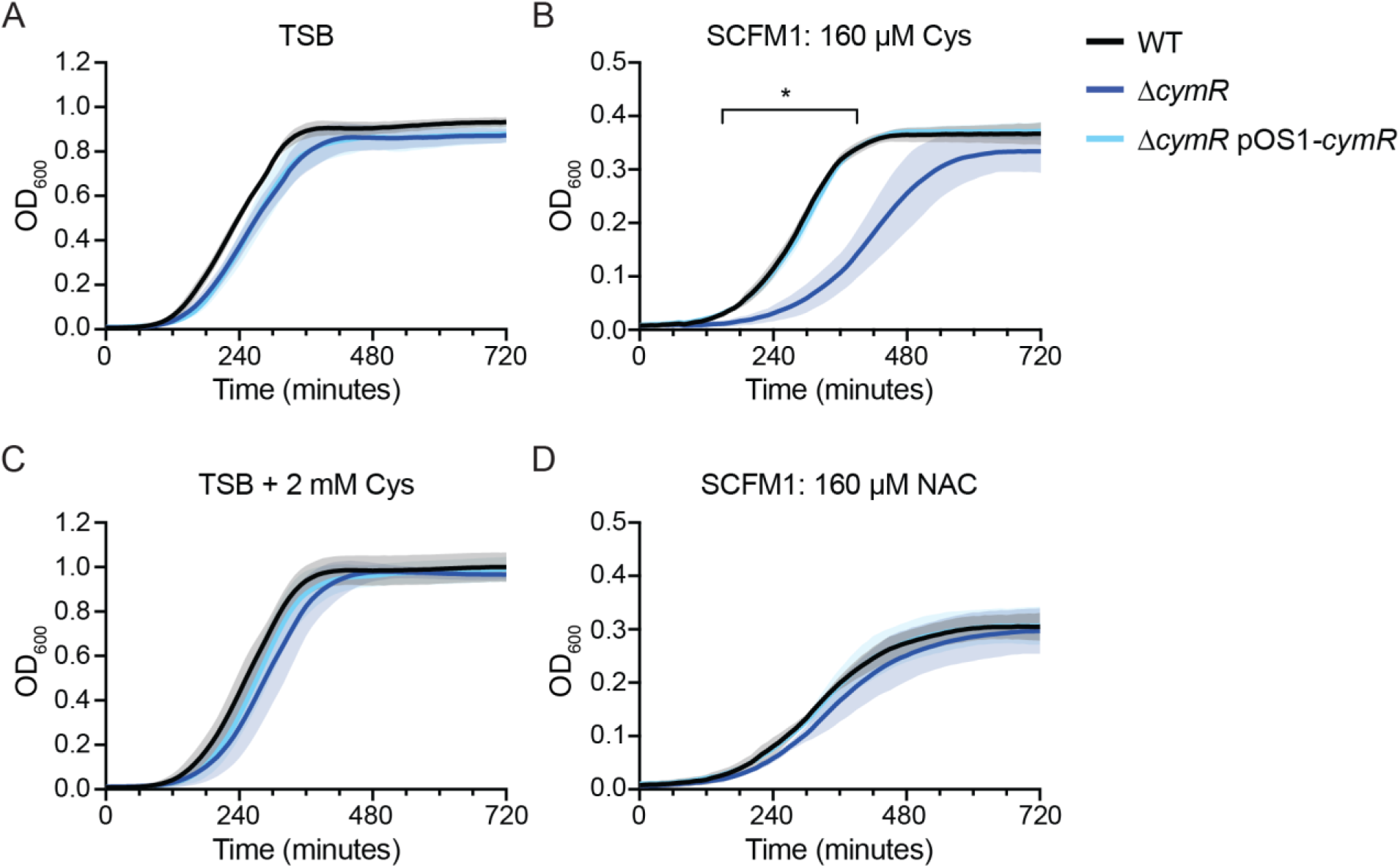
CymR-dependent growth kinetics in the presence of cysteine or N-**acetylcysteine**. Growth kinetics in **A)** TSB, **B)** SCFM1, **C)** TSB supplemented with 2 mM cysteine, and **D)** SCFM1 containing N-acetylcysteine (NAC) in place of cysteine. Data are the mean and standard deviation (SD) of three independent experiments. Asterisk (*) indicates *q* < 0.05 from 160 – 390 min, as measured by Welch’s T-test with two-stage step-up multiple comparison correction on OD_600_ measurements.

We then tested whether this growth defect was due to excess organic sulfur in general, or if it was specifically caused by cysteine. As *S. aureus* is a cysteine auxotroph, the cysteine in SCFM1 was replaced with the alternate organosulfur source N-acetylcysteine (NAC), which is imported by the same mechanism as cysteine (52). The Δ*cymR* mutant is able to grow similarly to WT in SCFM1 containing NAC in place of cysteine (Fig. 5D), suggesting that the growth defect of the Δ*cymR* mutant in SCFM1 is specifically due to cysteine.

Two of the direct repression targets of CymR are: *i) tcyABC*, encoding an ATP-binding cassette (ABC) transporter known to import cysteine, cystine, and NAC, and *ii) tcyP,* encoding a high-affinity permease that also imports cysteine, cystine, and NAC (52). As both *tcyABC* and *tcyP* are strongly upregulated in the absence of CymR, we hypothesized that uncontrolled cysteine import via TcyABC and/or TcyP was responsible for the growth defect of the Δ*cymR* mutant in SCFM1 (26). To test this hypothesis, we deleted *tcyA* and *tcyP* individually and in combination with deletion of Δ*cymR*. When grown in SCFM1, in which the only organic sulfur source is cysteine, the Δ*cymR* strain had a pronounced growth defect, as observed previously. Neither the Δ*tcyA* Δ*tcyP* double mutant nor the Δ*cymR* Δ*tcyA* Δ*tcyP* triple mutant were able to grow, supporting previous reports that TcyABC and TcyP are the only cysteine importers in *S. aureus* (Fig. 6A). Importantly, single Δ*tcyA* and Δ*tcyP* mutants grew similarly to WT in all conditions and deleting *tcyA* in the Δ*cymR* background partially rescued growth in the presence of cysteine (Fig. S2). When cysteine was substituted for equimolar organic sulfur in the form of GSH, which is imported independently of TcyABC and TcyP, all strains grew similarly to WT (Fig. 6B) (51, 52). In the presence of GSH to support the growth of mutants lacking cysteine importers, cysteine remained toxic to the Δ*cymR* mutant (Fig. 6C). Growth of Δ*cymR* was restored to WT levels when *tcyA* and *tcyP* were deleted in this background, even with the lowest concentration of GSH present in sputum (Fig. 6D). These results support the model that the Δ*cymR* mutant exhibits a growth defect in SCFM1 due to excess import of cysteine via TcyABC and TcyP.

**Figure 6.**
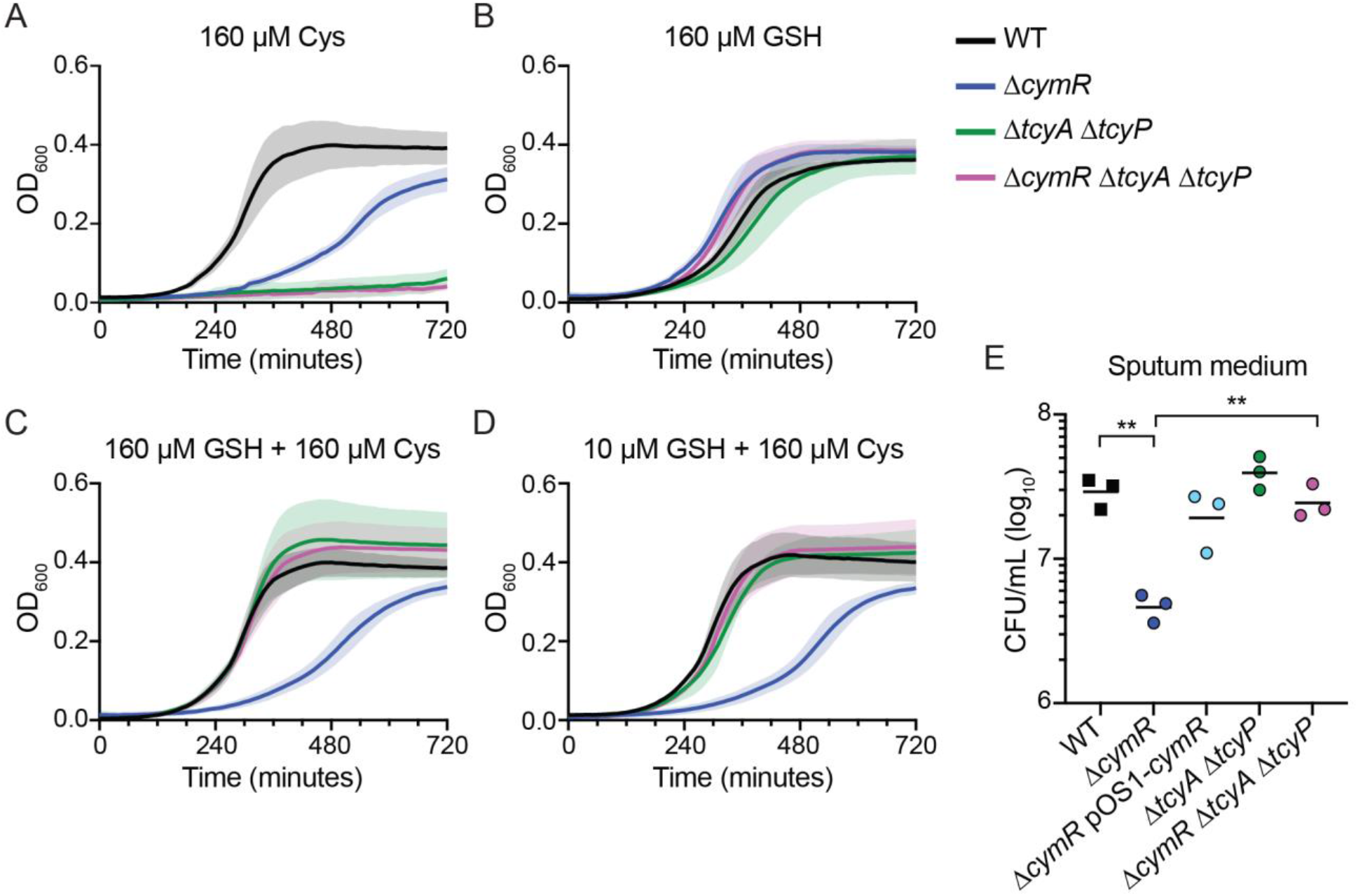
The role of cysteine importers in cysteine toxicity of Δ*cymR*. A-D) Growth kinetics of bacterial strains in SCFM1 containing the indicated sulfur source: A) cysteine, B) GSH, C) equimolar cysteine and GSH, D) cysteine and minimal GSH. Data are the mean and standard deviation (SD) of four independent experiments. **E)** Bacterial density 12 hours after inoculation in pooled sputum medium. Data from WT and Δ*cymR* strains are identical to those found in Figure 3. Two asterisks (**) indicate p < 0.01 as measured by Brown-Forsythe & Welch One-way ANOVA with Dunnett’s multiple comparison correction on log-transformed CFU values.

After determining the mechanism by which the Δ*cymR* mutant is attenuated for growth in the synthetic sputum model, we tested whether this same mechanism was responsible for the growth defect of the Δ*cymR* mutant in *ex vivo* CF sputum. We grew the Δ*tcyA* Δ*tcyP* double mutant and the Δ*cymR* Δ*tcyA* Δ*tcyP* triple mutant in the same pooled CF sputum medium that was used to test growth in monoculture for 12 hours. When *tcyA* and *tcyP* were deleted in the WT background, there was no difference in growth compared to WT (Fig. 6E). However, when both *tcyA* and *tcyP* were deleted in the Δ*cymR* background, growth was restored to WT levels. These data suggest that the Δ*cymR* mutant grows poorly in CF sputum due to excess cysteine import by TcyABC and TcyP.

## DISCUSSION

In this study we investigated the environment experienced by *S. aureus* in the CF lung, beginning with a Tn-seq screen to determine the *S. aureus* genes required in *ex vivo* CF sputum. By comparing four unique sputum samples as well as a pooled sputum medium, we identified 19 genes that were required for *S. aureus* growth in all sputum media and many more genes that were required for growth in at least one sputum medium. Mutants significantly depleted from all sputum samples include insertions in metabolic genes, those necessary for metal homeostasis, and the regulator of cysteine metabolism *cymR*. To decipher the roles of these genes in the unique environment of CF sputum, we quantified transition metals, the host protein calprotectin, and LMW thiols in sputum from 11 individuals. These analyses revealed that calprotectin was in stoichiometric excess relative to the total metals present, suggesting that bioavailable iron, copper, and zinc are substantially lower than the total measured concentrations (46–48). We also measured much more cysteine and GSH in sputum than previously appreciated. Together, results from this study provide a holistic understanding of the environment encountered by *S. aureus* during chronic infection of the CF lung and the genes it requires to survive and replicate in this milieu.

We performed Tn-seq in four unique media, each prepared from an independent CF sputum sample. This approach allowed us not only to identify genes that are generally required for growth in CF sputum, but to assess the extent to which the genetic requirements for growth are similar or different in distinct individuals. Previous studies, primarily in *P. aeruginosa*, used pooled CF sputa from multiple donors to minimize the effects of heterogeneity between samples. However, comparing Tn-seq results from individual sputum revealed phenotypes that were masked in the pooled medium, highlighting the power of exploring heterogeneity as a complement to studying generalizable phenotypes. For example, mutants of *graRS* and *vraGF* were enriched in two sputum media and depleted in two sputum media, while there was no major enrichment or depletion in the pooled medium. This phenotypic pattern suggests that there are important roles for *graRS* and *vraGF* during growth in CF sputum that likely differ from person-to-person.

Sputum heterogeneity may explain why some of the genes we selected for validation studies did not exhibit growth defects in monoculture, such as *nanE*, involved in the metabolism of sialic acid (29). Mucins are heavily decorated with sialic acid moieties and thus, sialic acid is abundant in CF sputum (53). Although *S. aureus* is unable to liberate sialic acid from mucin, other bacterial species commonly found in the CF lung, including *Schaalia odontolytica*, *Prevotella melaninongenica*, and *Streptococcus* spp., are capable of doing so (54–58). The requirement for *nanE* in some sputum media but not others could indicate sialic acid cross-feeding by other bacterial species present in those sputum samples.

One of the most prominent gene signatures we observed in the Tn-seq results was that of metal homeostasis. Mutants in genes involved in iron uptake and heme biosynthesis, including the *sir* and *fhu* operons as well as *hemL,* were significantly depleted in all four individual sputum media. Additionally, we identified the zinc-independent ribosomal protein encoded by *rpsN2* and the staphylopine exporter *cntE* as required for growth in sputum.

Together, these hits suggest that *S. aureus* experiences metal starvation in CF sputum. Somewhat paradoxically, we also identified genes important for detoxifying metal stress, including the manganese and copper exporters encoded by *mntE* and *copA*, respectively. Identifying multiple metal homeostasis genes prompted us to quantify the transition metals present in sputum.

Host sequestration of essential nutrient transition metals has long been appreciated to serve as a key defense against bacterial pathogens, a phenomenon known as nutritional immunity (38–40). Calprotectin is an abundant host protein that mediates nutritional immunity by binding and sequestering transition metals; thus, measuring calprotectin is critical for contextualizing metal availability (40, 46). Our analysis of calprotectin and nutrient metal concentrations in CF sputum revealed that calprotectin was far more abundant than copper, iron, or zinc in every sample tested. To our knowledge, sputum samples have not previously been analyzed for concentrations of both calprotectin and nutrient metals. ICP-MS quantifies all metals in a sample, whether or not they are protein-bound. Our data measuring the relative quantity of calprotectin suggests that the metals may not be accessible to bacteria.

The excess of calprotectin relative to nutrient metals in CF sputum suggests that Δ*cntE* and Δ*rpsN2* mutants exhibited significant growth defects in CF sputum medium due to metal starvation imposed by calprotectin. Loss of *cntE* prevents export of the metallophore staphylopine, which is primarily used for zinc acquisition (41). Sensitivity of Δ*cntE* to zinc starvation is at least partially due to toxic accumulation of staphylopine in the cytosol rather than the inability to acquire zinc (34, 35). To test whether the growth defect of the Δ*cntE* mutant is due to zinc starvation or staphylopine toxicity, we investigated the growth of a *cntA::Tn* mutant, which is impaired for staphylopine import (41). The mutant lacking *cntA* (*cntA*::Tn) exhibited an approximately 10-fold growth defect in CF sputum medium (*p* = 0.15). A possible explanation for this observation is that metal starvation plays a role in the growth defect of the Δ*cntE* mutant but that staphylopine toxicity may also contribute.

Recent studies have similarly underscored the selective pressure that calprotectin imposes on bacteria. For example, Tn-seq performed on *S. aureus* growing in TSB containing calprotectin identified genes that overlap strikingly with our CF sputum Tn-seq results, including the *cnt* operon and *rpsN2* (59). That is, simply adding calprotectin to rich medium is sufficient to recapitulate many phenotypes observed in CF sputum medium. Additionally, it was noted that adding calprotectin to SCFM improves the accuracy of SCFM in recapitulating the transcriptional landscape of *P. aeruginosa* during growth in CF sputum (60). The strong effect of calprotectin on *S. aureus* also likely explains why there was little overlap in our Tn-seq screen in SCFM3, which does not contain calprotectin, and our results in CF sputum. Our Tn-seq data are also interesting in the context of the *S. aureus* transcriptional response during growth in CF sputum. Many of the genes we identified as essential for growth in CF sputum media are also upregulated in *S. aureus* isolated from CF sputum samples, including the *cnt* operon, *nanE*, and *copA* (22). Ibberson and Whiteley mapped clinical isolate reads to a reduced core genome of 1,960 genes, so neither *rpsN2* nor *cymR* were analyzed; however, *tcyA* and *tcyP* were downregulated, consistent with our *cymR* analysis (22).

A Δ*cymR* deletion mutant exhibited a significant growth defect in CF sputum, which cannot be explained by calprotectin-mediated metal starvation and instead suggests a role for cysteine toxicity during growth in CF sputum. To explore whether cysteine could be present at toxic levels in CF sputum, we quantified the total amount of cysteine and GSH in the same samples in which we quantified calprotectin and metals. We found that cysteine concentrations ranged from 65 – 3,350 µM and GSH concentrations ranged from 11 - 296 µM, much higher quantities of both thiols than previously reported (20, 50). Excess cysteine has long been known to cause toxicity in bacteria, by either metabolic or oxidative stress (61–63). Our genetic data and thiol profiling results together suggest that in the absence of *cymR*, *S. aureus* likely imports a toxic amount of cysteine, causing attenuated growth in CF sputum. The exact mechanism by which cysteine is toxic to *S. aureus* in CF sputum is yet to be determined. The simplest explanation is that excess intracellular cysteine or oxidized cystine interacts with intracellular iron to inflict oxidative damage and attenuate growth, similar to what was shown in *E. coli* (63). However, we do not favor this hypothesis due to the fact that cysteine remained toxic to the Δ*cymR* mutant even in the presence of equimolar GSH, a potent antioxidant. Further, our thiol profiling data showed that GSH is present in CF sputum in high enough concentrations to fulfill the organic sulfur requirements of *S. aureus* without the need to import any free cysteine at all (51). An alternative hypothesis is that dysregulated cysteine import perturbs metabolic flux, leading to reduced growth. The dispensability of cysteine importers TcyABC and TcyP could suggest that *S. aureus* does not primarily use cysteine as a source of nutrient sulfur during growth in CF sputum as it does in other niches, including systemic infection (52). Avoiding free cysteine import altogether when other organosulfur sources are available could be a protective strategy against cysteine intoxication.

Although the manganese efflux pump *mntE* was identified by Tn-seq as important in all sputum media, the *mntE*::Tn mutant did not exhibit a growth defect in monoculture. This discrepancy is likely due to technical differences in the preparation of sputum medium. For Tn-seq experiments, sputum was homogenized in SMM, which contains manganese, while the sputum media used for validation studies were prepared in buffered base without manganese (20, 25). This hypothesis is supported by the fact that *mntE* was also identified as required for growth in SMM containing glucose in a pilot Tn-seq screen (data not shown). Additionally, the concentration of Mn (21 – 361 nM) we detected in sputum is not likely to cause toxicity (64, 65, 32). Mutants in the remaining two genes, *ylaN* and *pruA*, grew poorly compared to WT in monoculture but these growth defects were not statistically significant. This may be a result of the technical limitations of this growth model. WT *S. aureus* only replicates approximately 100-fold in sputum medium, so minor or variable growth defects are difficult to detect statistically, though they may be biologically relevant.

While we focused on genes required for growth in CF sputum, our Tn-seq screen also identified several mutants that were enriched in all four sputum media, indicating that these genes are actually detrimental to *S. aureus* growth in sputum. Most notably, transposon mutants in the *rsbUVW*-*sigB* operon were enriched across all of the sputum media. SigB is the general stress response sigma factor in *S. aureus*, and RsbU, RsbV, and RsbW are posttranslational regulators of SigB (66, 67). This finding is particularly interesting because this operon is known to accumulate inactivating mutations during long-term colonization of the CF lung (68). It has been hypothesized that isolates deficient in the general stress response emerge during long-term colonization due to their reduced virulence and, consequently, reduced immunogenicity (68). However, these genes were identified as hits in all four sputum media in the absence of an active immune response, suggesting that inactivation of the general stress response in *S. aureus* instead confers a general growth advantage in CF sputum.

In summary, we found that despite heterogeneity between samples, there exists a core set of genes required for growth in CF sputum. Many of these genes are involved in surviving metal starvation, which we propose is imposed by the high concentration of calprotectin present in CF sputum relative to available nutrient metals. We also establish that cysteine toxicity is a calprotectin-independent source of stress on *S. aureus* growing in CF sputum. These new insights into *S. aureus* growth in CF sputum fill an important gap in our understanding of how this key pathogen is able to establish and maintain infections in the CF lung environment.

## MATERIALS AND METHODS

### Bacterial strains and growth conditions

Bacterial strains used in this study are listed in Table S3. All *S. aureus* strains were cultivated in tryptic soy broth (TSB, BD Bacto) at 37 °C with aeration, unless otherwise indicated. All *E. coli* strains were grown in Miller’s lysogeny broth (LB) at 37 °C, with aeration. Plasmids were introduced into *E. coli* by chemical transformation and into *S. aureus* by electroporation. Antibiotics and additives (Sigma) were used at the following concentrations where appropriate: kanamycin, 50 µg/mL; chloramphenicol, 10 µg/mL (rich media) or 2.5 µg/mL (sputum media); erythromycin, 1 µg/mL; X-gal, 40 µg/mL. For all synthetic CF sputum medium (SCFM1 and SCFM3) experiments, SCFM was prepared as previously described with the exception that iron sulfate (FeSO_4_) was omitted to recapitulate observed metal scarcity in CF sputum (20, 21).

### Cloning and mutant construction

Plasmids, phages, and oligonucleotides used in this study are listed in Tables S4 and S5. Phage ɸ11:FRT was a gift from Timothy Meredith.(24) Plasmid pIMAY*-Z was constructed from pIMAY* and pMUTIN4. Plasmid pIMAY* was a gift from Angelika Gründling (Addgene plasmid #121441). Plasmid pMUTIN4 was obtained from the Bacillus Genetic Stock Center (BGSC; catalog #ECE139).

The Nebraska Transposon Mutant Library (NTML) was provided by the Network on Antimicrobial Resistance in *Staphylococcus aureus* (NARSA) for distribution through BEI Resources (NR-48501). Deletions were confirmed by PCR and Sanger sequencing. Transposon mutations sourced from the NTML were first confirmed by PCR and Sanger sequencing, then transduced into *S. aureus* JE2 using phage ɸ11::FRT, as described previously (23, 24, 69). Transductants were identified by antibiotic selection and confirmed by PCR and Sanger sequencing.

In-frame unmarked gene deletions in *S. aureus* were generated by allelic exchange using plasmid pIMAY*-Z and *E. coli* IM08B, as described previously (70–72). Complementation plasmids were generated by amplifying the coding sequence of the gene and its native promoter (approximately 200 bases preceding the coding sequence). Constructs were incorporated into shuttle vector pOS1 by restriction-ligation cloning before being introduced to *E. coli* IM08B by chemical transformation, followed by antibiotic selection. Inserts were confirmed by PCR and Sanger sequencing. Plasmid DNA was extracted from *E. coli* IM08B and introduced to *S. aureus* by electroporation and maintained by antibiotic selection.

### Sputum samples and sputum medium preparation

Sputum samples were donated by pediatric patients being treated for CF at Seattle Children’s Hospital and stored at -80 °C until processing. No patient data were associated with sputum samples for this study. To prepare growth media, frozen samples were thawed and homogenized by probe sonication at a concentration of 10% (w/v) in either Staphylococcal minimal medium with glucose and iron omitted (for Tn-seq) or SCFM buffered base (for CFU growth assays)(20, 25). Media not used immediately were stored at -80 °C until use. The ability of each medium to support growth of *S. aureus* JE2 was confirmed prior to inclusion of that medium in Tn-seq or growth assays.

### Transposon library construction, sequencing, and analysis

Plasmids pORF5-Tnp+, pORF5-Tnp-, and all transposon donor plasmids were a gift from Timothy Meredith (24). Transposon mutagenesis was carried out as previously described (24). Plasmids pORF5-Tnp+ (transposase-positive) and pORF5-Tnp-(transposase-negative control) were introduced to the target strain *S. aureus* JE2 by electroporation and maintained by antibiotic selection and growth at 30 °C. Transducing lysates were prepared with phage ɸ11::FRT from transposon donor strains carrying blunt, *P*_cap_, *P*_tuf_, *P*_erm_, and *P*_dual_ transposon variants. Each transposon construct was transduced into recipient strains JE2 pORF5-Tnp+ and JE2 pORF5-Tnp-, and transposition efficiency was confirmed by comparing the frequency of erythromycin-resistant colonies arising from the transposase-positive strain as compared to the transposase-negative control strain. For each construct in JE2 pORF5-Tnp+, approximately 10^5^ colonies were pooled, washed, and finally resuspended in TSB containing 20% glycerol for storage at -80 °C. The transposition process was completed two independent times for a total of ten libraries.

For Tn-seq, all libraries were thawed and pooled at equal volumes, then washed twice in phosphate-buffered saline (PBS) and resuspended in PBS to OD_600_ = 1.0 (approximately 10^9^ CFU/mL). Approximately 5 x 10^6^ CFU were used to inoculate 1 mL of media in each of the following 12 cultures: TSB; SCFM3; two cultures each of four individual SMM sputum media; and two cultures of SMM sputum medium composed of all four individual media pooled in equal volumes. Cultures were incubated at 37 °C with shaking at 200 rpm for 16 hours. CFU were enumerated by spot plating immediately after inoculation and 16 hours post-inoculation. After 16 hours of growth, 100 μL of each culture was used to inoculate 3 mL TSB. TSB subcultures were incubated at 37 °C with shaking at 200 rpm for eight hours to expand biomass. 1 mL aliquots of each TSB subculture were pelleted by centrifugation, the supernatant removed, and the cell pellet stored at -80 °C until DNA extraction.

Genomic DNA was isolated from cell pellets by phenol/chloroform extraction followed by ethanol precipitation. Libraries were prepared for sequencing as described previously (24). The resulting library was sequenced by Northwest Genomics Center using an Illumina NextSeq 500/550 platform with a PhiX spike-in of 40%. Sequencing results were demultiplexed using CutAdapt, aligned using bowtie2, and analyzed using TRANSIT (73–75). TRANSIT analysis parameters are included in Tables S1 and S2. Set analyses were visualized in R using the UpSetR package (76).

### Sputum growth assays

To quantify mutant growth in sputum, approximately 10^9^ bacterial cells were pelleted by centrifugation, washed in PBS, and resuspended in PBS at an OD_600_ of 0.1 (approximately 10^8^ CFU/mL). 2 µL (2 x 10^5^ CFU) of washed cells were used to inoculate 100 μL of TSB or sputum medium in a sterile polystyrene flat-bottom 96-well plate (Fisher). Plates were sealed with a sterile gas-permeable membrane (Diversified Biotech) and lid and incubated at 37 °C with shaking at 200 rpm for 12 hours. Immediately after inoculation and 12 hours post-inoculation, cultures were mixed by pipetting and CFU enumerated by dilution plating.

### Bacterial growth curve assays

Overnight bacterial cultures were washed in PBS and resuspended in PBS at an OD_600_ of 1.0. 2 μL of washed cells (2 x 10^6^ CFU) were used to inoculate 200 μL media in a flat-bottom 96-well plate with a gas-permeable membrane. Cysteine, glutathione, and N-acetylcysteine solutions were prepared immediately prior to each experiment. Plates were incubated in a BioTek Synergy HTX plate reader for 12 hours at 37 °C, with shaking, and growth was measured by OD_600_ every ten minutes for the duration of the experiment.

### Calprotectin quantification

Sputum (20% w/v) was treated with 0.1% DTT and mixed by pipetting or passing through a sterile 25g needle until homogeneous. An equal volume of PBS was added for a final sputum concentration of 10% (w/v). Debris was removed by centrifugation at 1200 rcf for 3 minutes followed by transfer of supernatant to a clean microcentrifuge tube. Calprotectin was quantified using a Human Calprotectin ELISA kit (Abcam). The assay was performed two independent times. Sputum supernatants were serially diluted in technical duplicate and all dilutions whose calprotectin concentrations fell within the range of the assay were used to compute the concentration of calprotectin in the undiluted sample. Calprotectin concentrations were converted to molarity by approximating the density of unprocessed sputum as 1 g/mL.

### Transition metal quantification

Transition metals were quantified by inductively coupled plasma mass spectrometry (ICP-MS). Samples were digested in metal-free 15 mL conical tubes in 200 µL 70% Optima-grade nitric acid at 65°C overnight, then diluted with UltraPure water to 20% nitric acid for analysis. Elemental quantification was conducted using an Agilent 7700 ICP-MS attached to an ASX-560 autosampler. The settings for analysis were cell entrance = −40 V, cell exit = −60 V, plate bias = −60 V, OctP bias = −18 V, and helium flow = 4.5 ml/min. Optimal voltages for extract 2, omega bias, omega lens, OctP RF, and deflect were empirically determined. Calibration curves for elements were generated using ARISTAR ICP standard mix. Samples were introduced by peristaltic pump with 0.5-mm-internal-diameter tubing through a MicroMist borosilicate glass nebulizer. They were initially taken up at 0.5 rps for 30 seconds, followed by 30 seconds at 0.1 rps to stabilize the signal. Spectrum mode analysis was performed at 0.1 rps, collecting three points across each peak and conducting three replicates of 100 sweeps for each element. The sampling probe and tubing were rinsed with 2% nitric acid for 30 seconds at 0.5 rps between each sample. Data were acquired and analyzed using Agilent MassHunter workstation software version A.01.02. Element quantities in parts per billion (ppb) were converted to molarity by approximating the density of unprocessed sputum as 1 g/mL.

### Low-molecular-weight thiol profiling

Sputum samples were resuspended to 10% sputum (w/v) in ultra-pure water. Samples were homogenized by sonication using a Sonic Dismembrator 550 (Fisher Scientific) with a microtip for one minute at maximum amplitude, 5.0 seconds on, 5.0 seconds off. To quantify total LMW thiol content, 100 µL of homogenized 10% sputum sample was mixed with TCEP in excess for a final concentration of 5 mM TCEP and left to react at RT for 5 minutes to reduce thiol disulfides. Samples were alkylated by addition of β-(4-hydroxyphenyl)ethyl iodoacetamide (HPE-IAM) (Chem-Impex, Catalog #23038) in excess, for a final concentration of 5 mM HPE-IAM in solution. Samples were incubated in a water bath at 37 °C for one hour. (77). Samples were clarified by centrifugation and the supernatant was filtered through a 0.22 µm spin filter (Fisher Scientific, Catalog #07200389). Samples were mixed in a 1:1 ratio of solution containing isotopically labeled standards of known concentration.

Alkylated standards were prepared using *D*_4_-HPE-IAM (Toronto Research Chemicals, Catalog #I685882) as described (78). Samples were analyzed by LC using C18 (YMC-Triart C18, 50 X 2.0 mm I.D., 1.9 µm) with an attached guard column (Phenomenex, UHPLC C18-Peptide, 2.1 mm I.D.), coupled to a Waters Synapt G2S mass spectrometer. Samples were run using mobile phase A (0.25% acetic acid, 10% methanol) and mobile phase B (0.25% acetic acid, 90% methanol) with the following LC elution gradient: 0-3 min, 100% A, 0% B; 3-7 min, linear gradient to 75% A, 25% B; 7-9 min, 75% A, 25% B; 9-12 min, linear gradient to 25% A, 75% B; 12-14 min, linear gradient to 0% A, 100% B; 14-20 min, 0% A, 100% B. The resulting total ion chromatogram (TIC) was searched for positively charged ions (*z* = 1; M^+^ or M + H^+^) (mass tolerance of ±0.02 Da) using Waters MassLynx software. The extracted ion chromatograms (EIC) of each light (H_4_) and heavy (*D*_4_) HPE-IAM-capped thiols identified in MS1 were obtained and peak areas were quantified. The ratio of light and heavy MS1 features (rang, 0.01 to 500) was used to calculate the concentration of each thiol using the known concentration of the heavy standard spiked into the mixture. All samples were run in technical triplicate and the data determined in units of pmol thiol/mg sputum. We then converted these data to µM thiol, assuming a value of 1 mL/g sputum, a value slightly higher than a partial specific volumes of protein (0.708 mL/g) or mucin (0.65 mL/g) (79).

## ACKNOWLEDGEMENTS

This work was supported by Cystic Fibrosis Foundation (CFF) awards RENIERE20I0 and SHULL22F0. Work in the Reniere lab is funded by NIH R01 AI132356. Work in the Hoffman lab is funded by CFF grants SINGH19R0 and HOFFMA20Y2-OUT. Work in the Skaar lab is funded by NIH R01s AI150701, AI1385, and AI073843. Work in the Giedroc lab is funded by NIH R35 GM118157. The funders had no role in study design, data collection and interpretation, or the decision to submit the work for publication. We thank Timothy Meredith for sharing ɸ11::FRT, bacterial strains, and protocols. Finally, we thank members of the CF community and their families for their participation in CF research, including this study.

## REFERENCES

1. Riordan JR, Rommens JM, Kerem B, Alon N, Rozmahel R, Grzelczak Z, Zielenski J, Lok S, Plavsic N, Chou JL. 1989. Identification of the cystic fibrosis gene: cloning and characterization of complementary DNA. Science 245:1066–1073.

2. Anderson MP, Gregory RJ, Thompson S, Souza DW, Paul S, Mulligan RC, Smith AE, Welsh MJ. 1991. Demonstration That CFTR Is a Chloride Channel by Alteration of Its Anion Selectivity. Science 253:202–205.

3. Bear CE, Li CH, Kartner N, Bridges RJ, Jensen TJ, Ramjeesingh M, Riordan JR. 1992. Purification and functional reconstitution of the cystic fibrosis transmembrane conductance regulator (CFTR). Cell 68:809–818.

4. Matsui H, Grubb BR, Tarran R, Randell SH, Gatzy JT, Davis CW, Boucher RC. 1998. Evidence for periciliary liquid layer depletion, not abnormal ion composition, in the pathogenesis of cystic fibrosis airways disease. Cell 95:1005–1015.

5. Boucher RC. 2007. Cystic fibrosis: a disease of vulnerability to airway surface dehydration. Trends in Molecular Medicine 13:231–240.

6. Ciofu O, Hansen CR, Høiby N. 2013. Respiratory bacterial infections in cystic fibrosis. Current Opinion in Pulmonary Medicine 19:251–258.

7. Cystic Fibrosis Foundation. 2022. Cystic Fibrosis Foundation Patient Registry 2022 Annual Data Report.

8. Dasenbrook EC, Merlo CA, Diener-West M, Lechtzin N, Boyle MP. 2008. Persistent methicillin-resistant Staphylococcus aureus and rate of FEV1 decline in cystic fibrosis. Am J Respir Crit Care Med 178:814–821.

9. Dasenbrook EC, Checkley W, Merlo CA, Konstan MW, Lechtzin N, Boyle MP. 2010. Association Between Respiratory Tract Methicillin-Resistant Staphylococcus aureus and Survival in Cystic Fibrosis. JAMA 303:2386–2392.

10. Vanderhelst E, De Meirleir L, Verbanck S, Piérard D, Vincken W, Malfroot A. 2012. Prevalence and impact on FEV(1) decline of chronic methicillin-resistant Staphylococcus aureus (MRSA) colonization in patients with cystic fibrosis. A single-center, case control study of 165 patients. J Cyst Fibros 11:2–7.

11. Esposito S, Pennoni G, Mencarini V, Palladino N, Peccini L, Principi N. 2019. Antimicrobial Treatment of Staphylococcus aureus in Patients With Cystic Fibrosis. Front Pharmacol 10:849.

12. Zampoli M, Morrow BM, Paul G. 2023. Real-world disparities and ethical considerations with access to CFTR modulator drugs: Mind the gap! Front Pharmacol 14:1163391.

13. Nichols DP, Morgan SJ, Skalland M, Vo AT, Van Dalfsen JM, Singh SB, Ni W, Hoffman LR, McGeer K, Heltshe SL, Clancy JP, Rowe SM, Jorth PK, Singh PK. 2023. Pharmacologic improvement of CFTR function rapidly decreases sputum pathogen density but lung infections generally persist. J Clin Invest e167957.

14. McCarron A, Parsons D, Donnelley M. 2021. Animal and Cell Culture Models for Cystic Fibrosis: Which Model Is Right for Your Application? The American Journal of Pathology 191:228–242.

15. Semaniakou A, Croll RP, Chappe V. 2019. Animal Models in the Pathophysiology of Cystic Fibrosis. Frontiers in Pharmacology 9:1475.

16. Stoltz DA, Meyerholz DK, Pezzulo AA, Ramachandran S, Rogan MP, Davis GJ, Hanfland RA, Wohlford-Lenane C, Dohrn CL, Bartlett JA, Nelson GA, Chang EH, Taft PJ, Ludwig PS, Estin M, Hornick EE, Launspach JL, Samuel M, Rokhlina T, Karp PH, Ostedgaard LS, Uc A, Starner TD, Horswill AR, Brogden KA, Prather RS, Richter SS, Shilyansky J, McCray PB, Zabner J, Welsh MJ. 2010. Cystic fibrosis pigs develop lung disease and exhibit defective bacterial eradication at birth. Sci Transl Med 2:29ra31.

17. Sun X, Sui H, Fisher JT, Yan Z, Liu X, Cho H-J, Joo NS, Zhang Y, Zhou W, Yi Y, Kinyon JM, Lei-Butters DC, Griffin MA, Naumann P, Luo M, Ascher J, Wang K, Frana T, Wine JJ, Meyerholz DK, Engelhardt JF. 2010. Disease phenotype of a ferret CFTR-knockout model of cystic fibrosis. J Clin Invest 120:3149–3160.

18. Van Goor F, Straley KS, Cao D, González J, Hadida S, Hazlewood A, Joubran J, Knapp T, Makings LR, Miller M, Neuberger T, Olson E, Panchenko V, Rader J, Singh A, Stack JH, Tung R, Grootenhuis PDJ, Negulescu P. 2006. Rescue of ΔF508-CFTR trafficking and gating in human cystic fibrosis airway primary cultures by small molecules. American Journal of Physiology-Lung Cellular and Molecular Physiology 290:L1117–L1130.

19. Ulrich M, Herbert S, Berger J, Bellon G, Louis D, Münker G, Döring G. 1998. Localization of Staphylococcus aureus in infected airways of patients with cystic fibrosis and in a cell culture model of S. aureus adherence. Am J Respir Cell Mol Biol 19:83–91.

20. Palmer KL, Aye LM, Whiteley M. 2007. Nutritional Cues Control Pseudomonas aeruginosa Multicellular Behavior in Cystic Fibrosis Sputum. Journal of Bacteriology 189:8079–8087.

21. Turner KH, Wessel AK, Palmer GC, Murray JL, Whiteley M. 2015. Essential genome of Pseudomonas aeruginosa in cystic fibrosis sputum. Proceedings of the National Academy of Sciences 112:4110–4115.

22. Ibberson CB, Whiteley M. 2019. The Staphylococcus aureus Transcriptome during Cystic Fibrosis Lung Infection. mBio 10:e02774–19.

23. Fey PD, Endres JL, Yajjala VK, Widhelm TJ, Boissy RJ, Bose JL, Bayles KW. 2013. A genetic resource for rapid and comprehensive phenotype screening of nonessential Staphylococcus aureus genes. mBio 4:e00537–00512.

24. Santiago M, Matano LM, Moussa SH, Gilmore MS, Walker S, Meredith TC. 2015. A new platform for ultra-high density Staphylococcus aureus transposon libraries. BMC Genomics 16:252.

25. Machado H, Weng LL, Dillon N, Seif Y, Holland M, Pekar JE, Monk JM, Nizet V, Palsson BO, Feist AM. 2019. Strain-Specific Metabolic Requirements Revealed by a Defined Minimal Medium for Systems Analyses of Staphylococcus aureus. Appl Environ Microbiol 85:e01773–19.

26. Soutourina O, Poupel O, Coppée J-Y, Danchin A, Msadek T, Martin-Verstraete I. 2009. CymR, the master regulator of cysteine metabolism in Staphylococcus aureus, controls host sulphur source utilization and plays a role in biofilm formation. Molecular Microbiology 73:194–211.

27. Soutourina O, Dubrac S, Poupel O, Msadek T, Martin-Verstraete I. 2010. The Pleiotropic CymR Regulator of Staphylococcus aureus Plays an Important Role in Virulence and Stress Response. PLOS Pathogens 6:e1000894.

28. Ji Q, Zhang L, Sun F, Deng X, Liang H, Bae T, He C. 2012. Staphylococcus aureus CymR Is a New Thiol-based Oxidation-sensing Regulator of Stress Resistance and Oxidative Response. J Biol Chem 287:21102–21109.

29. Olson ME, King JM, Yahr TL, Horswill AR. 2013. Sialic acid catabolism in Staphylococcus aureus. J Bacteriol 195:1779–1788.

30. Halsey CR, Lei S, Wax JK, Lehman MK, Nuxoll AS, Steinke L, Sadykov M, Powers R, Fey PD. 2017. Amino Acid Catabolism in Staphylococcus aureus and the Function of Carbon Catabolite Repression. mBio 8:10.1128/mbio.01434-16.

31. Ding X, Robbe-Masselot C, Fu X, Léonard R, Marsac B, Dauriat CJG, Lepissier A, Rytter H, Ramond E, Dupuis M, Euphrasie D, Dubail I, Schimmich C, Qin X, Parraga J, Leite-de-Moraes M, Ferroni A, Chassaing B, Sermet-Gaudelus I, Charbit A, Coureuil M, Jamet A. 2023. Airway environment drives the selection of quorum sensing mutants and promote Staphylococcus aureus chronic lifestyle. 1. Nat Commun 14:8135.

32. Grunenwald CM, Choby JE, Juttukonda LJ, Beavers WN, Weiss A, Torres VJ, Skaar EP. 2019. Manganese Detoxification by MntE Is Critical for Resistance to Oxidative Stress and Virulence of Staphylococcus aureus. mBio 10:e02915–18.

33. Sitthisak S, Knutsson L, Webb JW, Jayaswal RK. 2007. Molecular characterization of the copper transport system in Staphylococcus aureus. Microbiology 153:4274–4283.

34. Chen C, Hooper DC. 2019. Intracellular Accumulation of Staphylopine Impairs the Fitness of Staphylococcus aureus cntE Mutant. FEBS Lett 593:1213–1222.

35. Grim KP, Radin JN, Solórzano PKP, Morey JR, Frye KA, Ganio K, Neville SL, McDevitt CA, Kehl-Fie TE. 2020. Intracellular Accumulation of Staphylopine Can Sensitize Staphylococcus aureus to Host-Imposed Zinc Starvation by Chelation-Independent Toxicity. Journal of Bacteriology 202:10.1128/jb.00014-20.

36. Ghssein G, Brutesco C, Ouerdane L, Fojcik C, Izaute A, Wang S, Hajjar C, Lobinski R, Lemaire D, Richaud P, Voulhoux R, Espaillat A, Cava F, Pignol D, Borezée-Durant E, Arnoux P. 2016. Biosynthesis of a broad-spectrum nicotianamine-like metallophore in Staphylococcus aureus. Science 352:1105–1109.

37. Akanuma G, Kawamura F, Watanabe S, Watanabe M, Okawa F, Natori Y, Nanamiya H, Asai K, Chibazakura T, Yoshikawa H, Soma A, Hishida T, Kato-Yamada Y. 2021. Evolution of Ribosomal Protein S14 Demonstrated by the Reconstruction of Chimeric Ribosomes in Bacillus subtilis. J Bacteriol 203:e00599–20.

38. Kochan I. 1973. The Role of Iron in Bacterial Infections, with Special Consideration of Host-Tubercle Bacillus Interaction, p. 1–30. In Arber, W, Braun, W, Haas, R, Henle, W, Hofschneider, PH, Jerne, NK, Koldovský, P, Koprowski, H, Maaløe, O, Rott, R, Schweiger, HG, Sela, M, Syruček, L, Vogt, PK, Wecker, E (eds.), Current Topics in Microbiology and Immunology. Springer, Berlin, Heidelberg.

39. Weinberg ED. 1975. Nutritional Immunity: Host’s Attempt to Withhold Iron From Microbial Invaders. JAMA 231:39–41.

40. Hood MI, Skaar EP. 2012. Nutritional immunity: transition metals at the pathogen–host interface. Nat Rev Microbiol 10:525–537.

41. Grim KP, San Francisco B, Radin JN, Brazel EB, Kelliher JL, Párraga Solórzano PK, Kim PC, McDevitt CA, Kehl-Fie TE. 2017. The Metallophore Staphylopine Enables Staphylococcus aureus To Compete with the Host for Zinc and Overcome Nutritional Immunity. mBio 8:e01281–17.

42. Grim KP. 2020. Elucidating the mechanisms by which the human pathogen staphylococcus aureus resists host-imposed zinc starvation. Thesis. University of Illinois at Urbana-Champaign.

43. Boyd JM, Esquilín-Lebrón K, Campbell CJ, Kaler KR, Norambuena J, Foley ME, Stephens TG, Rios G, Mereddy G, Zheng V, Bovermann H, Kim J, Kulczyk AW, Yang JH, Greco TM, Cristea IM, Carabetta VJ, Beavers WN, Bhattacharya D, Skaar EP, Parker D, Carroll RK, Stemmler TL. 2023. YlaN is an iron(II) binding protein that functions to relieve Fur- mediated repression of gene expression in Staphylococcus aureus. bioRxiv 10.1101/2023.10.03.560778.

44. Sebulsky MT, Hohnstein D, Hunter MD, Heinrichs DE. 2000. Identification and characterization of a membrane permease involved in iron-hydroxamate transport in Staphylococcus aureus. J Bacteriol 182:4394–4400.

45. Dale SE, Sebulsky MT, Heinrichs DE. 2004. Involvement of SirABC in Iron-Siderophore Import in Staphylococcus aureus. Journal of Bacteriology 186:8356–8362.

46. Corbin BD, Seeley EH, Raab A, Feldmann J, Miller MR, Torres VJ, Anderson KL, Dattilo BM, Dunman PM, Gerads R, Caprioli RM, Nacken W, Chazin WJ, Skaar EP. 2008. Metal Chelation and Inhibition of Bacterial Growth in Tissue Abscesses. Science 319:962–965.

47. Nakashige TG, Zhang B, Krebs C, Nolan EM. 2015. Human Calprotectin Is an Iron- Sequestering Host-Defense Protein. Nat Chem Biol 11:765–771.

48. Besold AN, Gilston BA, Radin JN, Ramsoomair C, Culbertson EM, Li CX, Cormack BP, Chazin WJ, Kehl-Fie TE, Culotta VC. 2018. Role of Calprotectin in Withholding Zinc and Copper from Candida albicans. Infect Immun 86:e00779–17.

49. Gray RD, Imrie M, Boyd AC, Porteous D, Innes JA, Greening AP. 2010. Sputum and serum calprotectin are useful biomarkers during CF exacerbation. Journal of Cystic Fibrosis 9:193–198.

50. Dauletbaev N, Viel K, Buhl R, Wagner TOF, Bargon J. 2004. Glutathione and glutathione peroxidase in sputum samples of adult patients with cystic fibrosis. J Cyst Fibros 3:119– 124.

51. Lensmire JM, Wischer MR, Kraemer-Zimpel C, Kies PJ, Sosinski L, Ensink E, Dodson JP, Shook JC, Delekta PC, Cooper CC, Havlichek DH, Mulks MH, Lunt SY, Ravi J, Hammer ND. 2023. The glutathione import system satisfies the Staphylococcus aureus nutrient sulfur requirement and promotes interspecies competition. PLoS Genet 19:e1010834.

52. Lensmire JM, Dodson JP, Hsueh BY, Wischer MR, Delekta PC, Shook JC, Ottosen EN, Kies PJ, Ravi J, Hammer ND. 2020. The Staphylococcus aureus Cystine Transporters TcyABC and TcyP Facilitate Nutrient Sulfur Acquisition during Infection. Infect Immun 88:e00690–19.

53. Blix G, Lindberg E, Odin L, Werner I. 1955. Sialic Acids. Nature 175:340–341.

54. Ding X, Robbe-Masselot C, Fu X, Léonard R, Marsac B, Dauriat CJG, Lepissier A, Rytter H, Ramond E, Dupuis M, Euphrasie D, Dubail I, Schimmich C, Qin X, Parraga J, Leite-de- Moraes M, Ferroni A, Chassaing B, Sermet-Gaudelus I, Charbit A, Coureuil M, Jamet A. 2023. Airway environment drives the selection of quorum sensing mutants and promote Staphylococcus aureus chronic lifestyle. 1. Nat Commun 14:8135.

55. Byers HL, Tarelli E, Homer KA, Beighton D. 2000. Isolation and characterisation of sialidase from a strain of Streptococcus oralis. J Med Microbiol 49:235–244.

56. Beighton D, Whiley RA. 1990. Sialidase activity of the “Streptococcus milleri group” and other viridans group streptococci. J Clin Microbiol 28:1431–1433.

57. Moncla BJ, Braham P. 1989. Detection of sialidase (neuraminidase) activity in Actinomyces species by using 2’-(4-methylumbelliferyl)alpha-D-N-acetylneuraminic acid in a filter paper spot test. J Clin Microbiol 27:182–184.

58. Moncla BJ, Braham P, Rabe LK, Hillier SL. 1991. Rapid presumptive identification of black- pigmented gram-negative anaerobic bacteria by using 4-methylumbelliferone derivatives. J Clin Microbiol 29:1955–1958.

59. Reyes Ruiz VM, Freiberg JA, Weiss A, Green ER, Jobson M-E, Felton E, Shaw LN, Chazin WJ, Skaar EP. 2024. Coordinated adaptation of Staphylococcus aureus to calprotectin- dependent metal sequestration. mBio 0:e01389–24.

60. Lewin GR, Kapur A, Cornforth DM, Duncan RP, Diggle FL, Moustafa DA, Harrison SA, Skaar EP, Chazin WJ, Goldberg JB, Bomberger JM, Whiteley M. 2023. Application of a quantitative framework to improve the accuracy of a bacterial infection model. Proceedings of the National Academy of Sciences 120:e2221542120.

61. Sørensen MA, Pedersen S. 1991. Cysteine, even in low concentrations, induces transient amino acid starvation in Escherichia coli. Journal of Bacteriology 173:5244–5246.

62 . Kari C, Nagy Z, Kovács P, Hernádi F. 1971. Mechanism of the Growth Inhibitory Effect of Cysteine on Escherichia coli. Microbiology 68:349–356.

63. Park S, Imlay JA. 2003. High levels of intracellular cysteine promote oxidative DNA damage by driving the fenton reaction. J Bacteriol 185:1942–1950.

64. Kehl-Fie TE, Zhang Y, Moore JL, Farrand AJ, Hood MI, Rathi S, Chazin WJ, Caprioli RM, Skaar EP. 2013. MntABC and MntH Contribute to Systemic Staphylococcus aureus Infection by Competing with Calprotectin for Nutrient Manganese. Infect Immun 81:3395– 3405.

65. Handke LD, Gribenko AV, Timofeyeva Y, Scully IL, Anderson AS. 2018. MntC-Dependent Manganese Transport Is Essential for Staphylococcus aureus Oxidative Stress Resistance and Virulence. mSphere 3:10.1128/msphere.00336-18.

66. Pané-Farré J, Jonas B, Förstner K, Engelmann S, Hecker M. 2006. The σB regulon in Staphylococcus aureus and its regulation. International Journal of Medical Microbiology 296:237–258.

67. Pané-Farré J, Jonas B, Hardwick SW, Gronau K, Lewis RJ, Hecker M, Engelmann S. 2009. Role of RsbU in Controlling SigB Activity in Staphylococcus aureus following Alkaline Stress. J Bacteriol 191:2561–2573.

68. Long DR, Wolter DJ, Lee M, Precit M, McLean K, Holmes E, Penewit K, Waalkes A, Hoffman LR, Salipante SJ. 2021. Polyclonality, Shared Strains, and Convergent Evolution in Chronic Cystic Fibrosis Staphylococcus aureus Airway Infection. Am J Respir Crit Care Med 203:1127–1137.

69. Olson ME. 2016. Bacteriophage Transduction in Staphylococcus aureus, p. 69–74. In Bose, JL (ed.), The Genetic Manipulation of Staphylococci: Methods and Protocols. Springer, New York, NY.

70. Monk IR, Tree JJ, Howden BP, Stinear TP, Foster TJ. 2015. Complete Bypass of Restriction Systems for Major Staphylococcus aureus Lineages. mBio 6:e00308–00315.

71. Monk IR, Stinear TPY 2021. 2021. From cloning to mutant in 5 days: rapid allelic exchange in Staphylococcus aureus. Access Microbiology 3:000193.

72. Schuster CF, Howard SA, Gründling A 2019. 2019. Use of the counter selectable marker PheS* for genome engineering in Staphylococcus aureus. Microbiology 165:572–584.

73. Martin M. 2011. Cutadapt removes adapter sequences from high-throughput sequencing reads. 1. EMBnet.journal 17:10–12.

74. Langmead B, Salzberg SL. 2012. Fast gapped-read alignment with Bowtie 2. Nat Methods 9:357–359.

75. DeJesus MA, Ambadipudi C, Baker R, Sassetti C, Ioerger TR. 2015. TRANSIT - A Software Tool for Himar1 TnSeq Analysis. PLOS Computational Biology 11:e1004401.

76. Conway JR, Lex A, Gehlenborg N. 2017. UpSetR: an R package for the visualization of intersecting sets and their properties. Bioinformatics 33:2938–2940.

77. Zhang Y, Gonzalez-Gutierrez G, Legg KA, Walsh BJC, Pis Diez CM, Edmonds KA, Giedroc DP. 2022. Discovery and structure of a widespread bacterial ABC transporter specific for ergothioneine. Nat Commun 13:7586.

78. Abo M, Li C, Weerapana E. 2018. Isotopically-Labeled Iodoacetamide-Alkyne Probes for Quantitative Cysteine-Reactivity Profiling. Mol Pharmaceutics 15:743–749.

79. Thornton DJ, Sheehan JK, Lindgren H, Carlstedt I. 1991. Mucus glycoproteins from cystic fibrotic sputum. Macromolecular properties and structural “architecture”. Biochem J 276:667–675.

